# Developing a dengue forecast model using Long Short Term Memory neural networks method

**DOI:** 10.1101/760702

**Authors:** Jiucheng Xu, Keqiang Xu, Zhichao Li, Taotian Tu, Lei Xu, Qiyong Liu

## Abstract

**Background:** Dengue Fever (DF) is a tropical mosquito-borne disease that threatens public health and causes enormous economic burdens worldwide. In China, DF expanded from coastal region to inner land, and the incidence sharply increased in the last few years. In this study, we conduct the analysis of dengue using the Long Short Term Memory (LSTM) recurrent neural networks. This is an artificial intelligence technology, to develop a precise dengue forecast model.

**Methodology/Principal Findings:** The model is developed from monthly dengue cases and local meteorological data of 2005–2018 among top 20 Chinese cities with a record of the highest dengue incidence. The first 13 year data were used to construct the LSTM and to predict the dengue outbreaks in 2018. The results are compared with the estimated dengue cases of other previously published models. Model performance and prediction accuracy were assessed using Root Mean Square Error (RMSE). With the LSTM method, the prediction measurements of average RMSE drop by 54.79% and 34.76% as compared with the Susceptible Infected Recovered (SIR) model and Zero Inflated Generalized Additive Model (ZIGAM). Our results showed that if only local data were used to develop forecast models, the LSTM neural networks would fail to capture the transmission characteristics of dengue virus in areas with fewer dengue cases. Contrarily, transfer learning (TL) can improve the accuracy of prediction of the LSTM neural network model in areas with fewer dengue incidences.

**Conclusion and significance:** The LSTM model is beneficial in predicting dengue incidence as compared with other previously published forecasting models. The findings provide a more precise forecast dengue model, which can help the local government and health-related departments respond early to dengue epidemics.

**Author summary:** In China, DF is a public health concern that poses a great economic burden on local governments. However, the incidence has sharply increased in recent years with growth in the sub-regions. With this issue, it will be challenging to develop an accurate and timely dengue forecast model. LSTM recurrent neural networks, deep learning methods and virus propagation rules by learning from observational data offer more advantages in predicting the prevalence of infectious disease dynamics than the traditional statistical model. The 2005–2017 data of the top 20 Chinese cities with the highest dengue incidence were used to construct the LSTM model, advantageous in predicting dengue in most cities. Moreover, the model helped to predict the dengue outbreaks in 2018 and used to compare the estimated dengue cases with the RMSE results of other previously published models. A thorough search of the literature shows that this is the first established dengue forecast model using the LSTM method, which is effective in predicting the trend of dengue dynamics.

## Introduction

Dengue Fever (DF) has been a global threat since World War II [1,2], and each year, approximately 390 million dengue cases are reported worldwide, especially in Asia and South America [3,4]. The growing number of dengue cases is a huge economic burden [5]. In China, DF reemerged in Guangzhou city in 1978 after the city was free of the disease for more than 30 years [6]. Since then, more regions have reported dengue outbreaks, which include Guangdong, Guangxi, Yunnan and Zhejiang, and the incidence has increased steadily in recent years (Fig 1) [7]. Dengue was one of the most severe public health threats in China, especially when 45,230 dengue cases were reported in Guangdong Province in 2014 [8]. The administrative boundary data of China’s prefecture-level cities in 2005 and 2018 were derived from the National Geomatics Center of China [9].

**Fig. 1.**
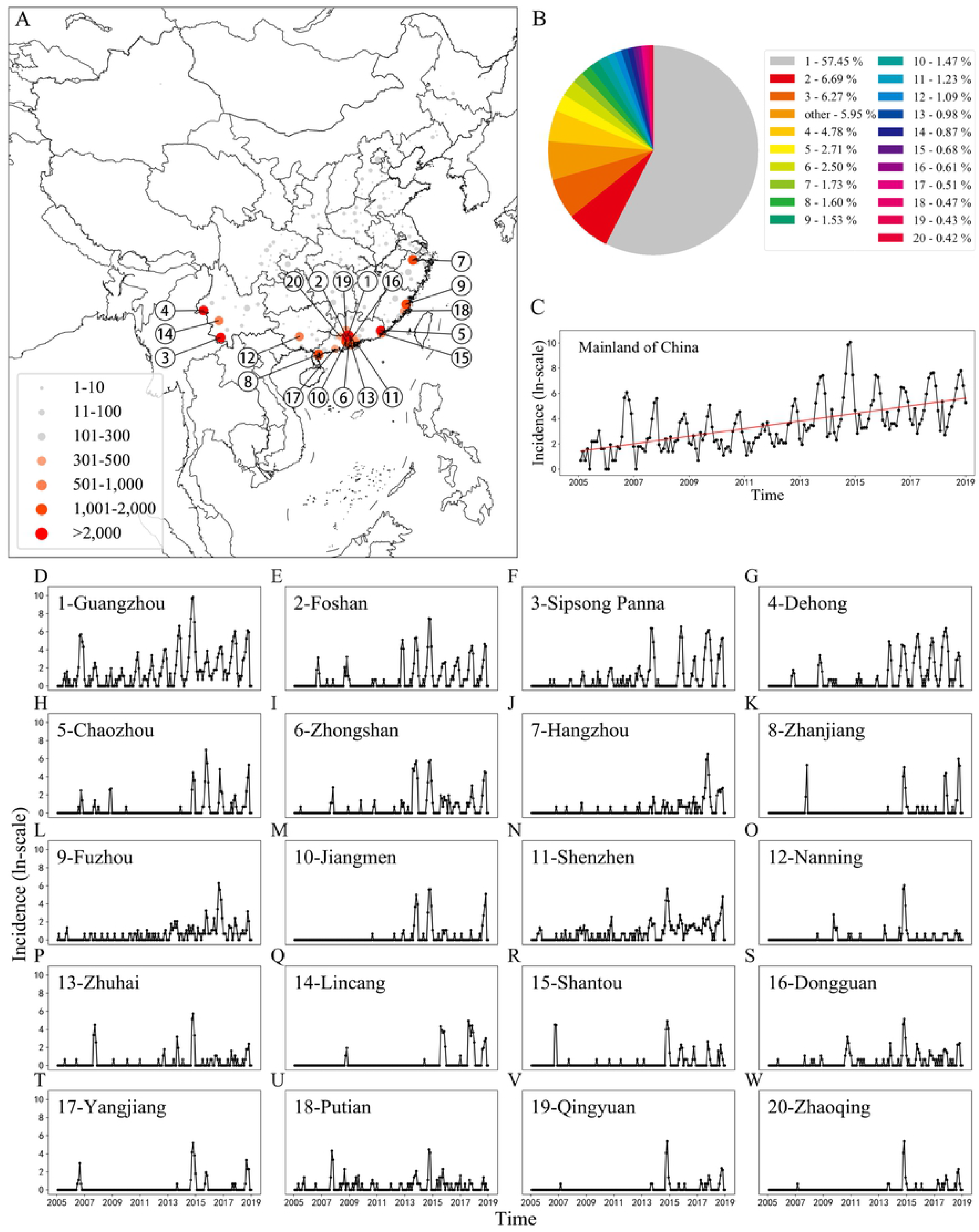
Spatial and temporal distribution of cases of dengue incidence in 2005–2018. (*A*) Distribution of dengue cases in China (case numbers and distinguished by color and size according to the magnitude in each city), (*B*) The proportion of cases in each city, (*C*) Times series of the dengue human incidence in Mainland China (on the logarithmic scale), (*D-W*) Times series of the dengue human incidence in the top 20 cities with the highest incidence of dengue (on the logarithmic scale).

DF is a mosquito-borne tropical disease caused by the dengue virus infection and a climate-sensitive disease [10,11]. The symptoms include high fever, headache, vomiting, muscle, joint pains and a skin rash [12]. In a small proportion, the disease develops into severe cases, resulting in bleeding, low levels of blood platelets and blood plasma leakage or shock syndrome [10]. The dengue virus (five serotypes of the dengue virus) is transmitted by the *Aedes albopictus* and *Aedes aegypti* [13], highly sensitive to environmental factors such as temperature and precipitation [14–17]. Meteorological conditions such as temperature, precipitation, and humidity have a significant impact on the DF spread as the conditions help to increase the *Aedes* population density [14–17]. As the temperature and the precipitation increases, the stages of development from *Aedes* larvae to pupa become shorter thereby increasing the *Aedes* population growth [18,19]. Conversely, high temperatures above 35°C or heavy precipitation may reduce the DF transmission by decreasing the *Aedes* survival rate [18,20]. Beside meteorological variables, social-economic factors also influence the spread of DF [21].

Early and accurate forecasts of dengue using different methods might minimise the threat, and this might help the government to take effective control measures. Numerous studies predicted the DF incidence using meteorological factors and vector population density. Shi et al. established a set of statistical models using the Least Absolute Shrinkage and Selection Operator method to improve forecasting of dengue in Singapore [22]. Xu et al. used Zero Inflated Generalized Additive Models (ZIGAM) to successfully demonstrate the effects of climatic conditions on the spread of mosquito and dengue transmission rate [8]. In addition, Li et al provide accurate dengue prediction by applying the Susceptible Infected Recovered (SIR) Epidemic Model in the mainland of China [23].

The Deep Learning method is a branch of Artificial Intelligence (AI) that generates virus propagation rules by learning from observational data. The method offers more advantages in predicting the prevalence of infectious disease dynamics as compared with the traditional statistical model [24,25]. Lee et al. showed that Artificial Neural Networks (ANNs) offer the potential benefit in forecasting fluctuations in mosquito population (especially the extreme values); this method is better than the traditional statistical techniques such as multiple regression model [26]. Aburas et al. predicted the number of dengue cases using ANNS, and the method produced encouraging results [27]. Truncating the gradient where this does not do harm using memory cells and gate units, LSTM recurrent neural networks can bridge a large amount of discrete-time steps [28]. Therefore, LSTM is one of the most advanced deep learning architectures for sequence learning tasks, such as speech recognition, or time series prediction [29,30]. LSTM has been used to forecast influenza trends and the epidemics of hand, foot and mouth disease successfully [31,32]. However, a thorough search of the literature shows that no previous attempt to deploy LSTM networks has been carried out to predict dengue incidence and assess its performance. In this study, we aimed to construct an accurate model to predict a monthly number of DF cases using the LSTM, a sophisticated deep learning method. This work addressed this gap a) by combining dengue monitoring data with meteorological data, proposing a prediction model for the DF incidence in China and b) by rigorously evaluating the predictive performance of the LSTM model and by comparing with different prediction algorithms.

## Methods

### Study Area

China is the most populous countries and the third-largest country in the world. A total of 250 cities in China reported dengue cases from 2005 to 2018. The top 20 cities with the highest incidence of dengue were selected as the study areas, highlighted in Fig 1A. However, Fig 1C demonstrated the number of confirmed trends and dengue cases in mainland China. This clearly showed a dengue outbreak in 2005 to 2018 confirmed cases of rising incidences. Fig 1B shows the proportion of the dengue cases in each city, and that of the Guangzhou city accounts for 57.45% of the total dengue cases in mainland China (Fig 1B). Fig 1(D-W) shows the time series of dengue cases in each city. From Fig 1D, the dengue epidemic in Guangzhou has obvious periodicity; with this feature, the DF data of Guangzhou were chosen to train the pre-training model.

### Dengue Incidence Data

Case-level records of human dengue incidence data from January 2005 to December 2018 were available in the China National Notifiable Disease Surveillance System. Relevant information for each case was recorded, including basic demographic characteristics (gender, age, nationality and residential address) and time of disease-related events (date of disease onset, diagnosis and death). All human dengue cases were diagnosed according to the diagnostic criteria for DF (WS216-2008) enacted by the Chinese Ministry of Health [33,34]. Human dengue cases per month (denoted as *D*) and monthly meteorological data were aggregated for the neural network model training and prediction.

### Meteorological Data and Attribute Selection

The climate data of these cities were obtained from the National Meteorological Information Center. A total of 15 meteorological variables was retained with no missing values in the raw data. These variables include three wind speed variables, four barometric variables, five temperature variables, two rainfall variables, and a humidity variable. The model will be overfitting if all variables are used for neural networks model training. Overfitting improves the model performances on the training set; however, it works poorly on the test set, revealing that the generalisation ability of the model is weak [35]. Thus, attribute selection is used to prevent overfitting and remove redundant attributes.

Attribute reduction for these meteorological variables is carried out using high correlation filtering and low variance filtering. Three meteorological variables were counted; these were the smallest variance in all the study areas and deleted. Four variables with the most frequent occurrence were deleted. Similarly, the two highest related meteorological variables were removed. Finally, nine weather conditions were retained through feature selection. The temperature variables include an average of daily highest temperature (denoted as *T^h^*), minimum air temperature (denoted as *T^min^*), and maximum air temperature (denoted as *T^max^*). Precipitation variables include an average of daily precipitation (denoted as *P^a^*), and a number of days with rainfall (denoted as *P^d^*). The pressure variables included maximum pressure (denoted a *Pr^max^*), an average of pressure (denoted as *Pr^a^*), and an average of water pressure (denoted as *Pr^w^*). The average of relative humidity is denoted as *H^a^*.

### Data set partition

The input data consists of the log-values of the human dengue cases and meteorological condition variables of the current month. The output data are the log-values of dengue cases in the subsequent month. The normalized data were divided into a training set, validation set and test set. The data from 2005 to 2016 were used as the training set, and the data of 2017 were used as the validation set to adjust network parameters such as the number of hidden. The data of 2018 were used as a test set. The main difference between train, validation and test sets is that the train set is used to adjust weight parameters of the neural networks and the validation set is used to assess the capabilities of the neural network model. However, the test set is the unseen data during the training. We calculated the performance index using the predicted value of the opposition number after the test was completed.

In this study, the dataset is relatively small for deep learning model; however, the climate has been proven as a driving force for DF [8,14]. This has a positive effect on the deep learning model, assisting in capturing the law of viral transmission and in predicting the number of cases.

### Model Rationale and Construction

LSTMs, as an advanced intelligent algorithm, can automatically find the characteristics of the long-term trend and the short-term fluctuation of time series data. LSTMs belong to the class of improved recurrent neural networks [28]. The ability of the LSTM neural network to find the connection automatically between attributes in time series is derived from learning. A learning process in the neural network model is a weight adjustment from giving examples, which makes the network output true observation without changes in the network structure [27]. LSTMs relies on the memory cell with three gating functions incorporated into its construction [36].

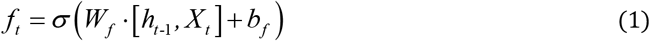

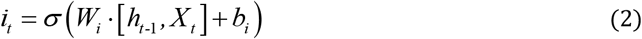

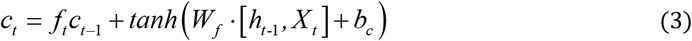

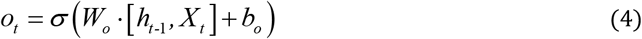

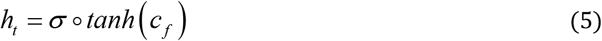

where *σ* is the logistic sigmoid function, and *f, i, c* and *o* are the *forget gate, input gate, cell* and *cell input* activation vectors and *output gate* respectively. *W* and *b* mean the weight matrix and the bias, respectively, and their subscripts have the obvious meaning. For example *W_f_, b_f_* are the weight matrix and the bias of the forget gate, whereas *W_i_, b_i_* are the weight matrix and the bias of the input gate.

The LSTM memory cell was used to compose network prediction architecture (Fig 2). The 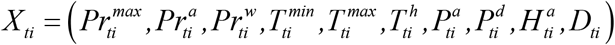 is a set of input vector sequence at the same month. The *c* = (*c*_1_, *c*_2_, *c*_64_) is the number of the cell of hidden layers. Input time series *X* = (*X*_*t*1_, *X*_*t*2_,*X*_*t*12_) is transmitted to the hidden layers, that contain n memory cell in everyone, through weighted connections to compute output *Y*, the incidence of dengue in the subsequent month.

**Fig. 2.**
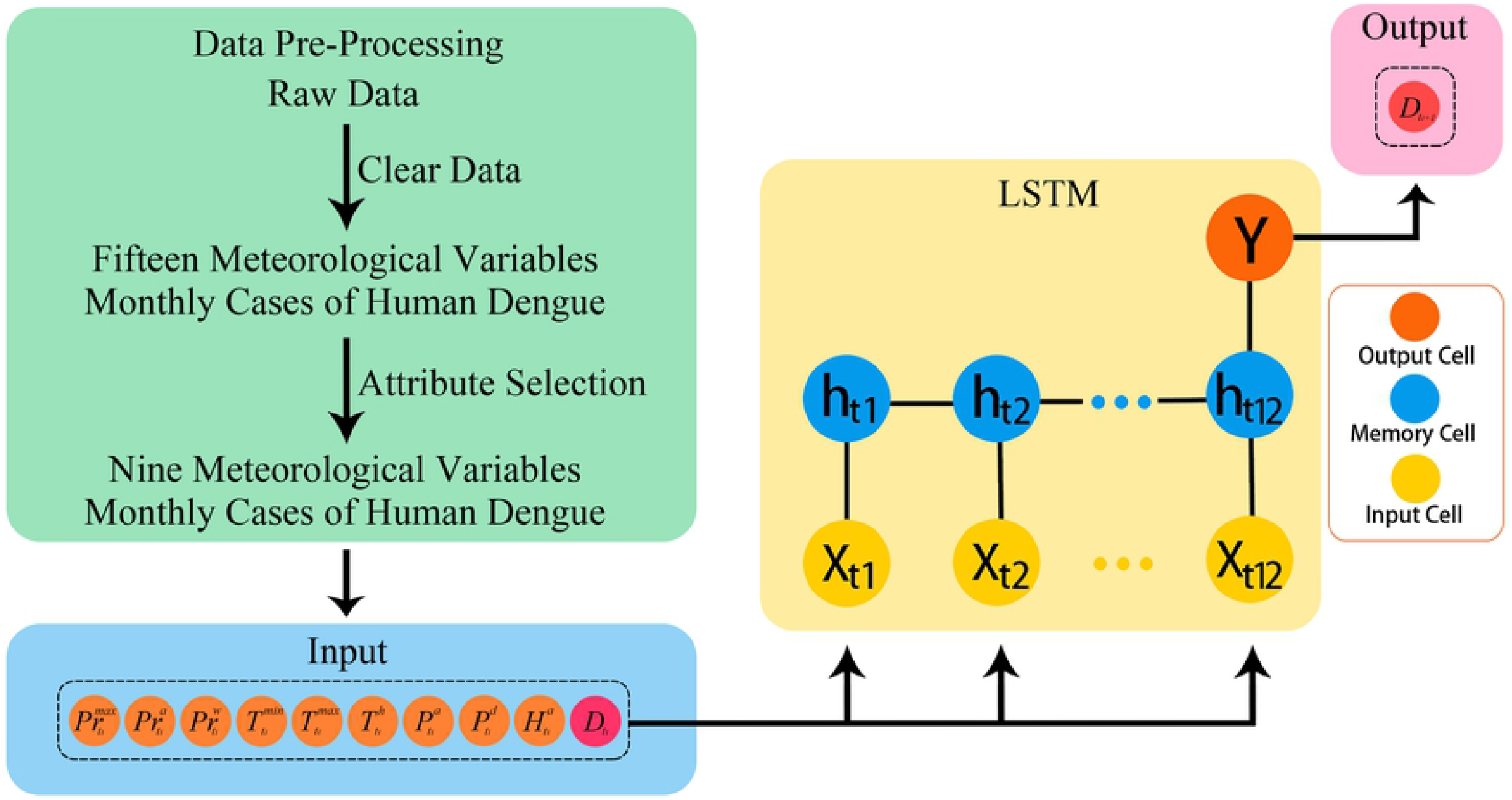
The architecture of the dengue forecast model using LSTMs network.

The hidden layer activations are calculated by iterating through the following equations from *t*1 to *t*12:

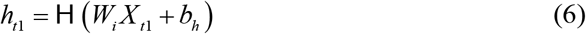

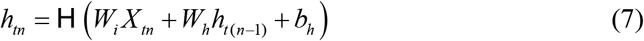

where the *W* terms represent the weight matrices, the *b_h_* is bias vector of the hidden layer and H is the function of the hidden layer.

The output is computed through the following equations:

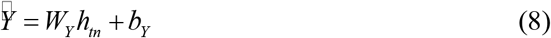

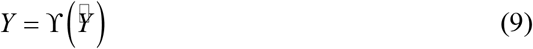

where *W_Y_* is the weight matrix of fully connected layer, the *b_Y_* is a bias vector of output layer and ϒ is the output layer function.

## Results

### Final parameter selection

The LSTM network model has three layers: a hidden layer which includes 64 memory cells, an input layer and an output layer (Fig. 2). The LSTM model consisted of 10 input parameters such as monthly mean maximum temperature, monthly average relative humidity, monthly raining days and the observation taken last month. These parameters are represented with 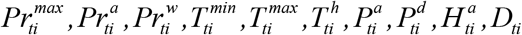 in Fig 2. Since the transmission cycle of dengue virus is one year in China, the time step of the LSTM network is set to 12. The result in the validation set was optimum when the number of the hidden layer was set to one (Table 1). The model need to must input 12 sets of parameters to produce an output the number of dengue cases in the next month. These inputs are shown in *X*_*t*1_ to *X*_*t*12_ and the output is shown in *Y* in Fig 2.

**Table 1.** RMSE validation set for a different number of hidden layers in LSTM model.

| Hidden Layer<br>($i$) | RMSE |
| --- | --- |
| $i = 1$ | **21.81** |
| $i = 2$ | 31.22 |
| $i = 3$ | 132.57 |

### Model training

The internal weight parameters of the LSTM neural network were adjusted through Adam optimizer. The initial learning rate of the model was 1*e*^-4^, and the training time was 8000 epochs. The dropout rate was set to 40% to combat network overfitting. All models were trained in python using the TensorFlow library.

We designed two model training routes, one is to train the LSTM model using the local data only; the second is to train a pre-train model using data from Guangzhou and train models of other cities using transfer learning (TL). Moreover, the Guangzhou data were selected to train the pre-training model as they contained a large number of DF cases. This is the LSTM model that learns the concepts of mapping the input (Meteorological Data and Dengue Incidence Data) and output data (Dengue Incidence Data in next month). The model fit of the Guangzhou data is the starting point for a model of the second city.

### Model evaluation

Model performance and prediction accuracy were measured by RMSE [37]. RMSE is widely used to evaluate continuous variables by measuring the differences between predicted and observed values.

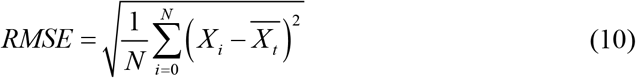

Fig 3 presented the prediction results of network models using different training strategies. The TL (red lines) produced less RMSE in most cities used to select the model of TL technology to predict the human incidence.

**Fig. 3.**
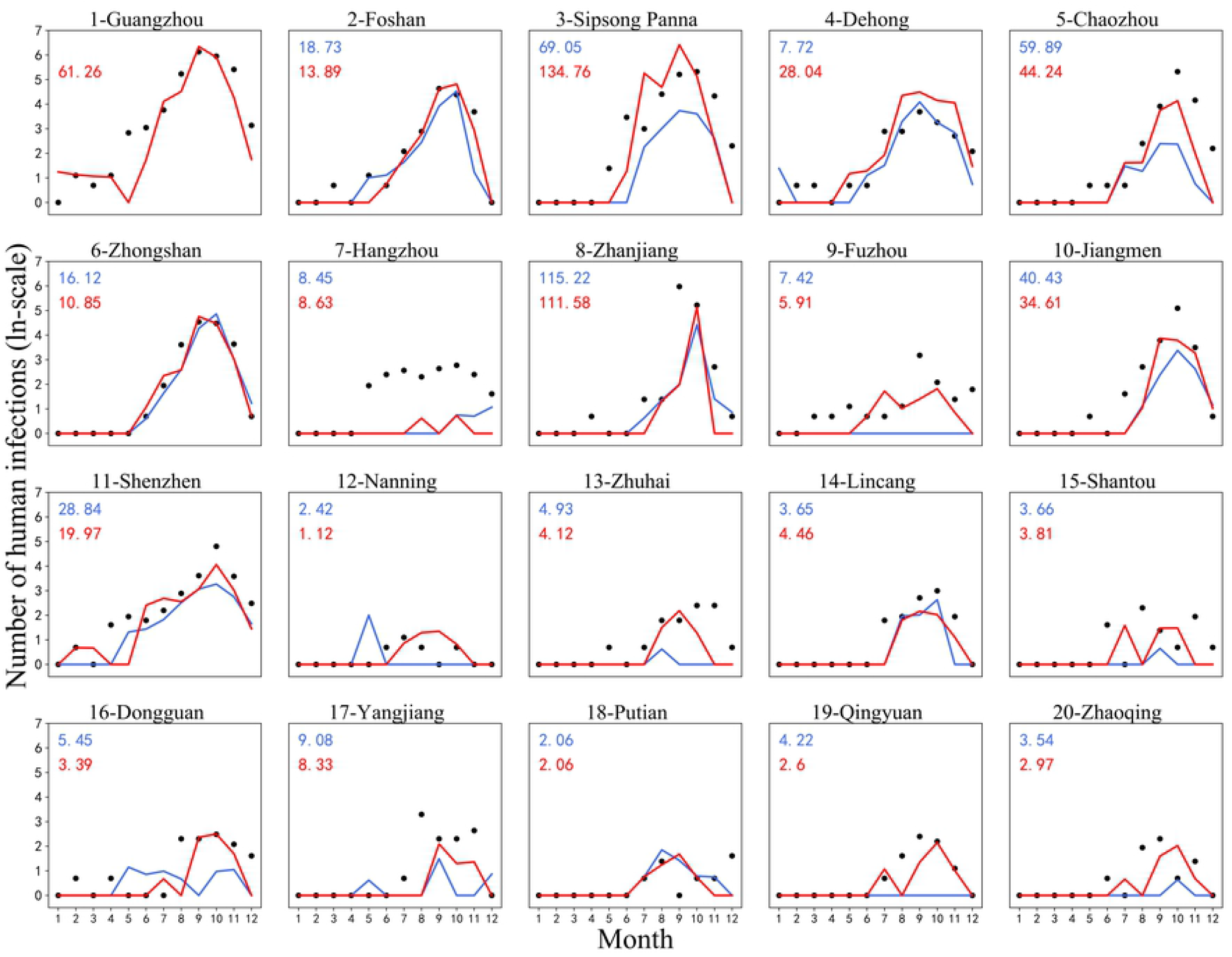
Diagnosed and predicted human cases of dengue across various cities in 2018. The diagnosed number of human cases in 2005 – 2017 was used for model training and selection. The LSTM was used TL prediction of human cases (red lines), and LSTM learning prediction of human cases (blue lines) were compared with observed data (black point) on the logarithmic scale.

## Discussion

This study demonstrates an efficient tool using the LSTM neural network, an AI algorithm, to predict dengue outbreaks in mainland China. Our results imply that the LSTM method has the potential obtaining epidemic trends of dengue than the SIR and ZIGAM models. Notably, the LSTM method can make the average RMSE prediction measurements drop by 54.79% and 34.76% as compared with the SIR model and ZIGAM model (Table 2). Moreover, the LSTM algorithm can be applied to more regions because the deep learning algorithm learns the transmission characteristics of the dengue virus propagation directly from the original data (Table 2). Thus, this paper uses only the health department case monitoring data and meteorological data to develop a DF prediction model based on the LSTM recursive neural network without mosquito density data.

**Table 2.** Comparison of model performance and goodness-of-fít using the RMSE.

| City | Human Incidence | LSTMs | LSTMs Transfer Learning | SIR | ZIGAM | SVR |
| --- | --- | --- | --- | --- | --- | --- |
| 1-Guangzhou | 42,764 | 61.26 | - | 49.17 | 42.52 | 96.96 |
| 2-Foshan | 4,986 | 18.73 | **13.89** | 16.89 | 86.33 | 16.01 |
| 3-Sipsong Panna | 4,770 | **69.05** | 134.76 | - | - | - |
| 4-Dehong | 3,542 | **7.72** | 28.04 | - | - | - |
| 5-Chaozhou | 2,108 | 59.89 | **44.24** | - | - | - |
| 6-Zhongshan | 1,935 | 16.12 | 10.85 | 43.20 | 6.35 | 1.10 |
| 7-Hangzhou | 1,287 | **8.45** | 8.63 | - | - | - |
| 8-Zhanjiang | 1,218 | 115.22 | **111.58** | - | - | - |
| 9-Fuzhou | 1,163 | 7.42 | **5.91** | - | - | - |
| 10-Jiangmen | 1,138 | 40.43 | **34.61** | - | - | - |
| 11-Shenzhen | 994 | 28.84 | 19.97 | 30.76 | 27.83 | 1.64 |
| 12-Nanning | 801 | 2.42 | **1.12** | - | - | - |
| 13-Zhuhai | 740 | 4.93 | 4.12 | 35.36 | 18.24 | 1.39 |
| 14Lincang | 655 | **3.65** | 4.46 | - | - | - |
| 15-Shantou | 511 | **3.66** | 3.81 | 64.30 | 13.05 | - |
| 16-Dongguan | 461 | 5.45 | **3.39** | - | - | - |
| 17-Yangjiang | 395 | 9.08 | **8.33** | - | - | - |
| 18-Putian | 355 | 2.06 | **2.06** | 93.03 | 64.03 | - |
| 19-Qingyuan | 353 | 4.22 | **2.60** | - | - | - |
| 20-Zhaoqing | 316 | 3.54 | **2.97** | - | - | - |

Monthly dengue cases and meteorological data were chosen to develop the LSTM model without mosquito density data. Dengue cases data and meteorological data can be updated promptly in relevant departments. However, the timely update of the local mosquito data cannot be done in China. Previous studies [8,14] reveal that the relationship between dengue dynamics and climatic variation can be understood. In this study, mosquito data were discarded to assist the LSTM model to predict the prevalence of dengue.

Our neural network model is compared with the other three previously published models. Table 2 presented the comparison of different models to forecast the dengue outbreak in the city of China. The prediction error of the Guangdong of LSTM model is slightly higher than that of the SIR and ZIGAM models, owing to the removal of mosquito data. However, in terms of prediction accuracy, the LSTM neural network model has a low RMSE in most cities than other models. With the application of the LSTM model to more regions in mainland China, it was found that the LSTM failed to capture the characteristics of viral transmission in areas with a low dengue incidence. However, this problem can be solved using TL in areas shown in Table 2.

Using TL, the RMSE prediction results of some models significant declined. In these areas, RMSE decline is more obvious in the vicinity of Guangzhou. The neural networks model is less effective for low-resource training. Although TL improves the model appropriately, it is required that the source and target tasks are the same [38]. The TL is an optimisation to improve the learning of a new task through the transfer of knowledge from related learning tasks. In our study, TL is available in similar climate regious, and it is the improvement of learning in areas with fewer dengue incidences through the transfer of the model to already learned areas with high dengue incidences.

Our study develops statistical analysis though the multiyear time series [8,23,27], built from the previous research that links dengue with meteorological variables [14–17] and climate change [39]. Our findings suggest that the advance prediction of dengue trends is achieved through neural network models using combinations of meteorological and disease surveillance variables.

However, our modelling approaches have some limitations. The RMSE of the predicted result is high in Hangzhou, where no record of dengue outbreaks is found. Moreover, we failed to consider potential relevant other social factors, such as population movements and city size. In the future, we will obtain a more accurate prediction model by obtaining traffic and immigration data. Nevertheless, this study provides a more efficient technique for the prediction of infectious diseases.

## Acknowledgements

We are grateful to all units involved the collection of DF data and infectious disease surveillance systems, coupled with the funders of this study. This research was partially supported by the donations from Delos Living LLC and the Cyrus Tang Foundation to Tsinghua University. This work was supported by the National Natural Science Foundation of China (NFSC) (Nos. 61772176, 61370169, 61402153, 41801336 and 61976082), the Plan for Scientific Innovation Talent of Henan Province (No. 184100510003), the Science and Technology Department, Henan Province (Nos. 182102210362 and 162102210261), the Young Scholar Program of Henan Province (No. 2017GGJS041), the Key Scientific and Technological Project of Xinxiang City of China (No. CXGG17002).

